# Using eDNA to Detect the Distribution and Density of Invasive Crayfish in the Honghe-Hani Rice Terrace World Heritage Site

**DOI:** 10.1101/109074

**Authors:** Wang Cai, Zhuxin Ma, Chunyan Yang, Lin Wang, Wenzhi Wang, Guigang Zhao, Yupeng Geng, Douglas W. Yu

## Abstract

The Honghe-Hani landscape in China is a UNESCO World Natural Heritage site due to the beauty of its thousands of rice terraces, but these structures are in danger from the invasive crayfish *Procambarus clarkii*. Crayfish dig nest holes, which collapse terrace walls and destroy rice production. Under the current control strategy, farmers self-report crayfish and are issued pesticide, but this strategy is not expected to eradicate the crayfish nor to prevent their spread since farmers are not able to detect small numbers of crayfish. Thus, we tested whether environmental DNA (eDNA) from paddy-water samples could provide a sensitive detection method. In an aquarium experiment, Real-time Quantitative polymerase chain reaction (qPCR) successfully detected crayfish, even at a simulated density of one crayfish per average-sized paddy (with one false negative). In a field test, we tested eDNA and bottle traps against direct counts of crayfish. eDNA successfully detected crayfish in all 25 paddies where crayfish were observed and in none of the 7 paddies where crayfish were absent. Bottle-trapping was successful in only 68% of the crayfish-present paddies. eDNA concentrations also correlated positively with crayfish counts. In sum, these results suggest that single samples of eDNA are able to detect small crayfish populations, but not perfectly. Thus, we conclude that a program of repeated eDNA sampling is now feasible and likely reliable for measuring crayfish geographic range and for detecting new invasion fronts in the Honghe Hani landscape, which would inform regional control efforts and help to prevent the further spread of this invasive crayfish.

## Introduction

Invasive species are generally recognized as having severe negative effects on native species and ecosystems [1]. Being able to detect small populations would help achieve local or regional eradication [2-3]. The Honghe Hani Rice Terrace landscape in Yuanyang, Yunnan, China is a UNESCO World Natural Heritage site (whc.unesco.org/en/list/1111/, accessed 21 Nov 2016) due to the beauty of its thousands of terraced rice paddies (Fig 1). However, as of 2011, 29,501 ha of rice paddies have been reported to be occupied by the invasive crayfish *Procambarus clarkii* (Cambaridae, Decapoda), which was introduced in 2006 by a local farmer who wanted to breed them for sale. A local newspaper has reported that farmers are sometimes able to catch over 1,000 crayfish in a single rice paddy of average size (= 0.1 ha) [4]. According to the report, crayfish populations dig multiple nest holes, causing water leakage and collapsed walls, thereby destroying the terraced landscape and rice production. In 2013, the crayfish population was deemed widespread [4].

**Fig 1.**
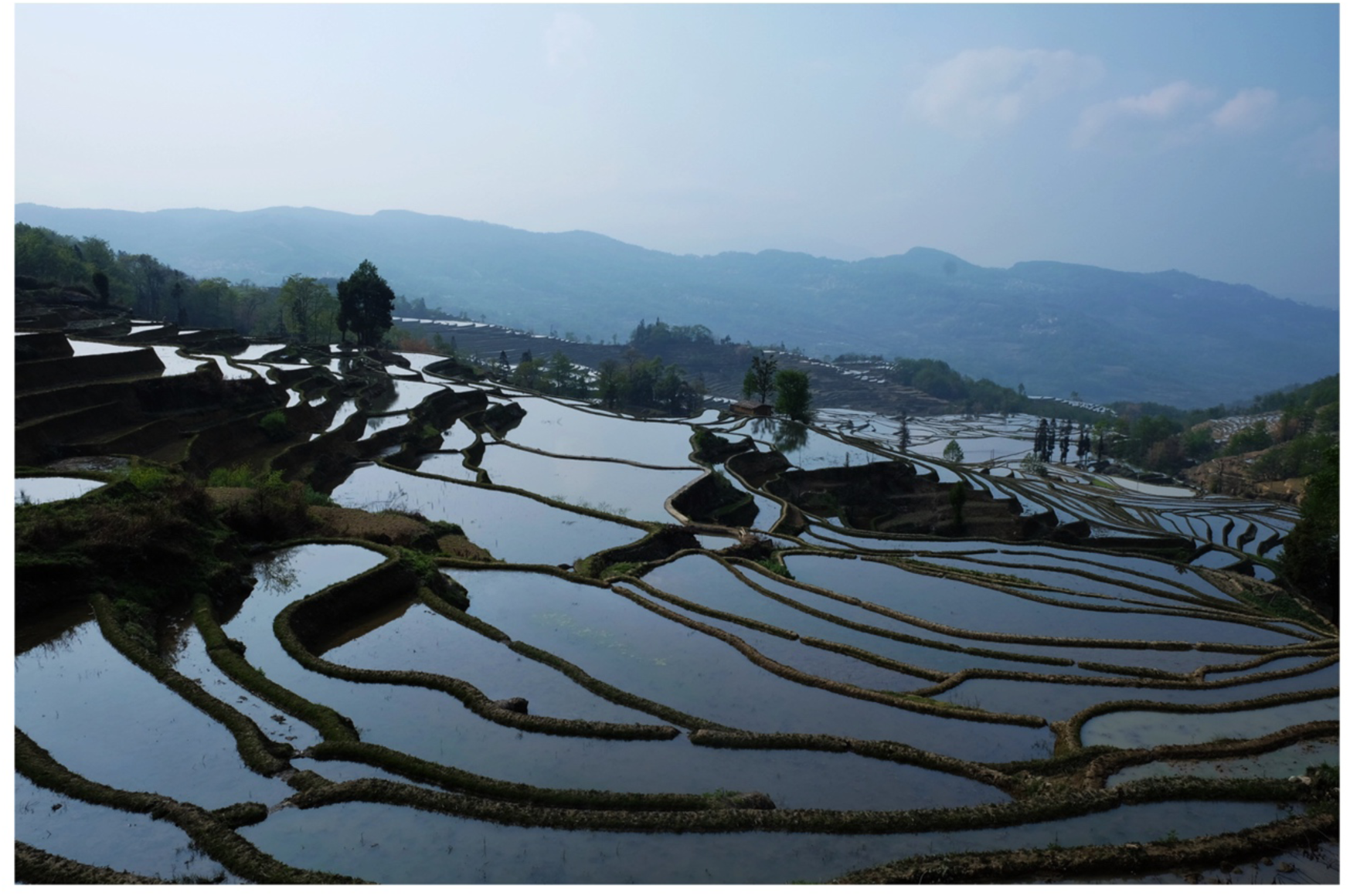
**Honghe Hani Terrace landscape in Yunnan, China,** 30 March 2016.

The current control strategy relies exclusively on farmer self-reporting after the detection of crayfish damage to a rice paddy. The farmers are issued deltamethrin pesticide, for which the local government spends 1.1 million RMB (~US$162,000 in November 2016) every year. The pesticide is applied in two seasons: before the transplantation of rice seedlings into the terrace in March and after the rice harvest in October. This strategy only kills crayfish in the focal paddy (which typically is at high density) and therefore has not resulted in the eradication of the crayfish population, nor is it expected to, because farmers can easily fail to detect *P. clarkii* crayfish at low numbers (false negatives), which can happen at the early stages of an invasion and also when local populations have not been fully eradicated. It is therefore necessary to be able to detect low densities of crayfish, which would help to prevent new populations from establishing and also to ensure that full local eradication takes place [5]. Conversely, farmers might falsely or incorrectly report crayfish presence (false positives), leading to over-application of pesticide.

Bottle-trap-sampling is the conventional approach for detecting invasive crayfish [6]. However, bottle-trap-sampling is inefficient because it requires at least two site visits (to set out and to collect) and, as we will show, because it often fails to detect crayfish.

Environmental DNA (eDNA) is a rapidly growing monitoring method that has been used for invasive species detection and management in aquatic systems [7-8]. eDNA refers to genetic material that is extracted from bulk environmental samples such as water, soil, and air [9-10], derived from shed tissue by the target species. eDNA is considered more sensitive than traditional techniques for detecting aquatic invaders [11-12]. Furthermore, eDNA has been used for estimating population abundances and thus might be a useful tool for aquatic invasive species detection in the initial dispersal stage [13-14].

Most previous aquatic-eDNA studies have focused on fish and amphibians, but two studies have used eDNA to detect freshwater crayfish [15-16]. Unlike fish and amphibians, which release abundant extracellular DNA via body mucus secretion [17], crayfish and other crustacean species seem to release limited amounts of tissue into water, which makes them more difficult to detect.

In this study, we tested the use of eDNA from rice-paddy water to detect *P. clarkii*. First, we used an aquarium dilution experiment to test if eDNA concentration correlates with crayfish density, which is known to be challenging [18-21], and to estimate the minimum number of crayfish that is likely to be detectable in an average (0.1-ha) rice paddy. Second, we compared the efficiencies of farmer reports, eDNA, and bottle-trap-sampling in the field. Finally, we mapped crayfish detections across a landscape gradient of high to low density.

## Materials and methods

### Study area

Field samples were collected in the Honghe Hani Terraces, Yunnan province, southwest China (23.1 °N, 102.8 °W). This region is suitable for growing rice due to the annual average temperature of 15 °C and abundant sunshine. Most terrace slopes are between 15-75 degrees, extending from the floor at 150 meters above sea level to more than 2000 masl. Our study was conducted during 15 to 30 March, 2016.

### Aquarium experiment

We used an aquarium experiment to test the extent to which qPCR of water samples can detect crayfish at low simulated densities. Crayfish-free water samples from rice paddies for the aquarium experiment were collected in the villages of Bada, Quanfuzhuang, and Duoyishu, where crayfish have not been reported by farmers, found in traps, nor using qPCR. We collected 60 L of paddy water in each of the three locations in March 2016, transported the water to Kunming city, and used it to create three aquaria respectively in which we reared *P. clarkii* crayfish (purchased at a market), after using qPCR to confirm the absence of crayfish DNA (authors’ unpubl. data). Paddy water was used in the aquaria to simulate the field environment as closely as possible, including possible qPCR interference.

Each of the three aquaria consisted of a polyethylene tank (60 × 40 × 10 cm) in an isolated location to prevent cross contamination. All tanks were first sterilized via 10-min exposure to a 10% bleach solution and then rinsed thoroughly before rearing crayfish. Water sieve and filter apparatus were originally cleaned by high-pressure steam sterilization and packed separately. We also created one crayfish-free control aquarium.

At the start of the experiment, adult *P. clarkii* crayfish (male, mean length 9 cm, mean biomass 12 g) were reared at a water temperature of 17±2 °C under a natural summer photoperiod (10 hours daylight), which mimics the field environment. Each tank contained 3 crayfish and 3 L of paddy water (1 crayfish / L) during rearing. The crayfish were fed 10 g of pork liver once per day at 8 am. Crayfish that died or were visibly injured from fighting were replaced as soon as detected (n = 4), to avoid crayfish releasing extra eDNA that could skew results.

After three days, we removed all crayfish and carried out a dilution series for each aquarium. The beginning density was one crayfish / L, and for each dilution step, we mixed 1-L of aquarium water with 9-L of the original crayfish-free paddy water in a new tank. We created seven densities: one crayfish per 1, 10, 10^2^, 10^3^, 10^4^, 10^5^, and 10^6^ L. The 1 crayfish/10^5^ L density is equivalent to 1 crayfish / average paddy, because the average paddy contains 1×10^5^ L water (0.1 ha at an average water depth of 0.1 m), under the assumption that all released eDNA first floats into the water (but see Discussion below regarding this assumption). Twenty-two water samples (seven densities X three aquaria plus one crayfish-free control aquarium) were then taken and filtered for eDNA, as follows.

From each aquarium, we used a 500-mL sterile bottle to remove a 1-L total water sample 5 cm below the surface at 5 locations evenly distributed around the aquarium, which was pooled in a disposable plastic bottle. We then used a metal sieve (pore size = 20 μm, diameter = 10 cm) to remove solid impurities because they block filter pores and might preserve long-term DNA [22-23], whereas we are looking for fresh eDNA. We previously determined that this sieve minimized eDNA loss while removing most sediment (authors’ unpubl. data).

After sieving, we filtered 1 L of each water sample through a disposable filter funnel (MicroFunnel™ST, PALL, Michigan, US) containing a 47-mm diameter cellulose nitrate filter paper with 0.45 μm pore size. Both a peristaltic pump and a hand pump were used in our sampling. We handled and stored the filter papers in the same way for each sample following the method outlined in [24]. The filter paper was rolled up using a new and previously autoclaved forceps and placed into a 2 mL tube with 900 μL absolute ethanol. Each tube was sealed and placed in an individually labeled plastic bag. Samples were stored at room temperature until DNA extraction (to mimic field transport conditions), which was conducted within two weeks.

### Field sampling and surveys

We compared four detection methods for crayfish: (1) farmer interviews, (2) collection of water samples for eDNA, (3), bottle-trap-sampling, and (4) directly counting the crayfish in each surveyed paddy, taking advantage of farmer application of pesticide during our surveys. Thirty-two rice paddies were sampled in the villages of Qingkou (n_rice paddies_ = 18), Duoyishu (n = 8), and Duoyishuxiao (n = 6), where crayfish have been reported.

At each sampling site, we asked the resident farmer if crayfish were currently present in the paddy and to judge subjectively if the crayfish density was high, medium, or low. Farmers estimated the crayfish density based on direct observation, whether they had used pesticides to eliminate crayfish over the last few years, and physical evidence of terrace damage. We did not attempt to normalize this judgment across farmers.

We collected paddy water for eDNA, using a protocol adapted from Treguier [15] and Pilliod [25]. In short, at each paddy, we used a 500-mL sterile bottle to collect a 1-L total water sample from 5 cm below the surface at 15 locations evenly distributed throughout the paddy, pooling the subsamples into a single disposable plastic bottle. We then sieved, filtered, and preserved samples following the *aquarium experiment*, and transported the samples to Kunming city at room temperature until DNA extraction, which was conducted within two weeks. One L of distilled water was used as a negative control and was filtered in the field after collecting the field samples, using the field apparatus. We used this negative control to detect possible sources of contamination from the apparatus, filter-paper handling procedures, and subsequent laboratory procedures.

We also used bottle-trap-sampling to catch crayfish, which is the conventional method for crayfish detection, using 3, 5, 7, or 9 traps, respectively, in paddies of 0-200, 200-500, 500-1000, or > 1000 m^2^, following Treguier [15]. Each trap was a collapsible cylinder [26] (35 cm × 17 cm ×17 cm) of polyamide wire (5 mm mesh) with two side entrances (with an inner opening diameter of 5 cm), and baited with 10 g of pork liver. Traps were evenly placed in the morning (8:00-10:00 am) on the bottom of the paddy, after the water sampling, and retrieved 24 hours later, following Dougherty [16], at which point we counted any crayfish and released non-target organisms.

Following all sampling, we directly counted the crayfish in each paddy, by taking advantage of the fact that farmers in Yuanyang annually kill crayfish with pesticide before planting rice seedlings. Crayfish hide in nest holes, but applying pesticide forces crayfish to leave their holes. In our study paddies, farmers applied pesticide after we sampled the water and removed the bottle traps. Twenty-four hours later, the water level was lowered by draining, and we counted all crayfish in the paddy. We selected paddies from 7 to 500 m^2^, which were small enough for us to count all crayfish in them, and then normalized the counts to a density for the average paddy of 0.1 Ha (10^5^ L water).

### Ethics statement

All of the rice paddies were sampled from 15 to 30 March 2016, with the landowners’ permission. The pesticide was legally applied by the landowners. The dosage of pesticide followed the product instruction manual and information provided by the local agricultural agent. After counting, farmers disposed of the captured crayfish.

### eDNA extraction and qPCR

The eDNA samples were extracted from the filters. Filters were first taken out from the ethanol solution and allowed to air dry for 12 hours. DNA extraction used the DNeasy Blood and Tissue Kit (Qiagen, Hilden, Germany) following the manufacturer’s protocol, except that we used 567 μL of buffer ATL and 63 μL of proteinase K [27], and the tubes were incubated at 56 °C for 48 h to allow for a more complete lysis [28]. We also eluted samples with 100 μL of TE buffer twice (200 μL total).

An extra DNA sample extract from *P. clarkii* tissue was used as a positive control, quantified using a NanoDrop (Thermo Fisher Scientific, Inc., Waltham, MA, USA), and added to every qPCR plate. We also created a dilution series with this sample (10^-1^ to 10^-9^ ng/μL) to estimate the Limit Of Quantification (LOQ, the lowest concentration of target DNA that can yield an acceptable level of precision and accuracy) and the Limit Of Detection (LOD, the minimum concentration of target DNA that can be detected in the sample).

Taqman qPCR assays used the *SPY_ProCla_F* and *SPY_ProCla_R* primers and the *SPY_ProCla* probe [15], with the following reagents: 3.6 μL template DNA, 10 μL TaqMan Environmental Master Mix 2.0 (Life Technologies, Carlsbad, California, USA), 4 μL ddH_2_O, 0.8 μL of each primer (10 μM) and 0.8 μL of probe (2.5 μM). Each eDNA sample was run in three technical replicates (for LOD and LOQ testing, we used ten replicates) on a QuantStudio 12K Flex Real-Time PCR System (Life Technologies). Three negative controls were added to every PCR plate: (1) double-distilled H_2_O filtered in the field or crayfish-free water from the aquarium, depending on sample source, (2) double-distilled H_2_O from the extraction process, and (3) double-distilled H_2_O from the qPCR planning. qPCR thermal cycling was carried out as follows: 50 °C for 5 min and 95 °C for 10 min, followed by 55 cycles of 95 °C for 30 s, and 56 °C for 1 min.

Samples that were not detected by qPCR could be false negatives (crayfish present but eDNA not detected because of inhibition or concentration below the LOD) or true negatives (crayfish not detected because they were indeed absent). To test for inhibition, we added crayfish tissue DNA at a concentration of 10^-5^ ng/μl, which is above the LOD (see below), to the negative samples and re-ran the qPCR.

### Statistical analyses

CCT is a conversion value of the threshold cycle of qPCR detection (CCT = 1000 / threshold cycle, cT), where cT is the ‘critical threshold’ number of qPCR cycles required for positive detection. We use CCT for visualization because CCT rises with eDNA concentration. Crayfish densities in the aquarium experiment were from the dilution gradient (1 crayfish / L to 1 crayfish / 10^6^ L). Crayfish densities in field samples were from the counts normalized to an estimated number of crayfish per average paddy of 10^5^ L (6 crayfish / average paddy to 2,000 crayfish / average paddy). The bottle-trap-sampling index is the number of crayfish caught per trap per paddy.

We compared the two estimates using a linear regression with two starting predictors: CCT and bottle-trap-sampling index, which is conventionally applied as an index of relative abundance of crayfish [29, 30]. Analyses were performed using *R* 3.1.3 (R Core Team, 2013).

## Results

### Limits of detection and the quantification experiment

Using *P. clarkii* tissue, the Limit Of Quantification (LOQ) was estimated to be 10^-4^ ng_DNA_/μL, with CCT_mean_ = 34.01, and the Limit Of Detection (LOD) was estimated to be 10^-8^ ng_DNA_/μL with CCT_mean_ = 23.05, which is similar to Treguier *et al*.’s results [15] (Fig 2).

**Fig 2.**
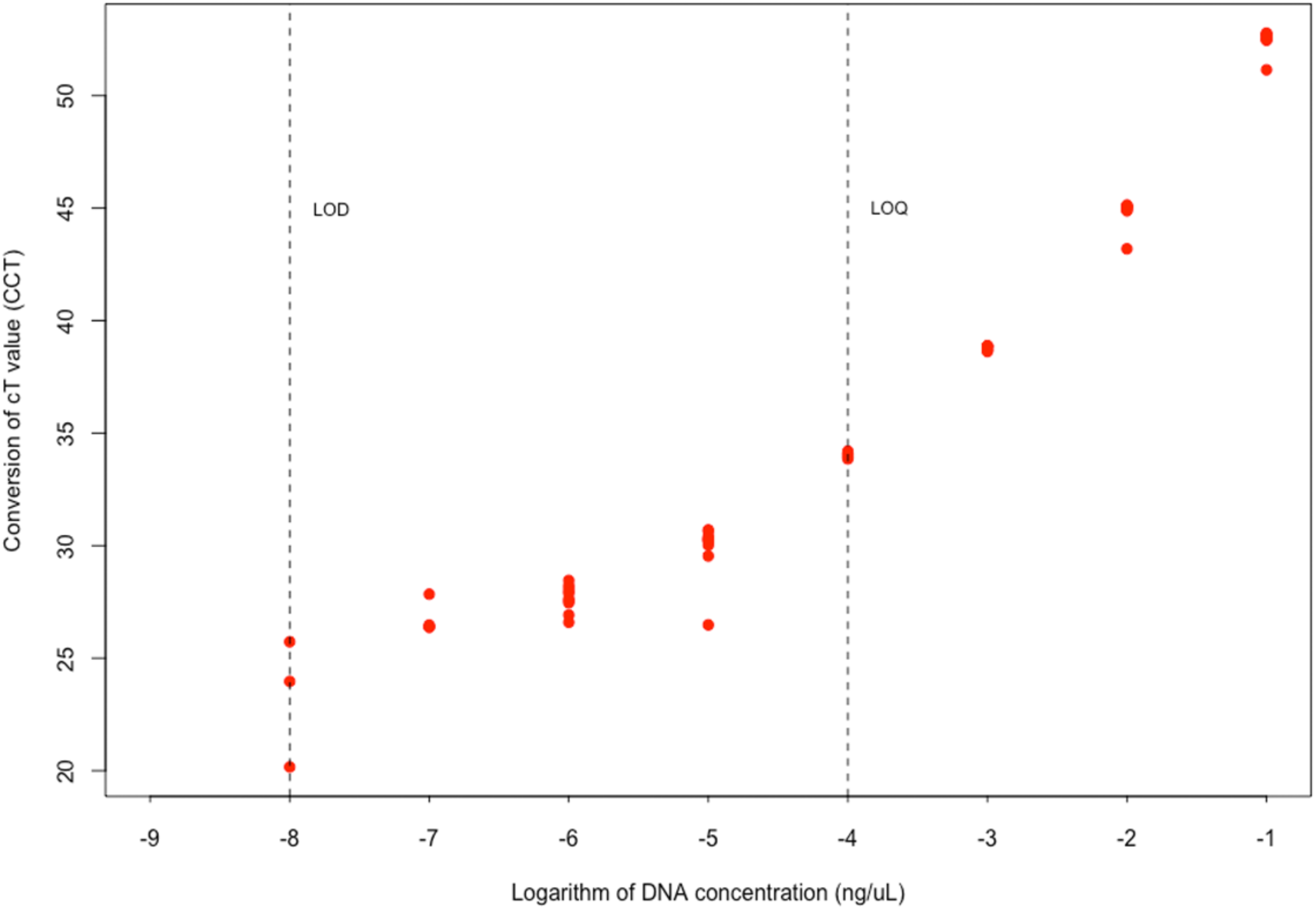
qPCR limit of quantification (LOQ) and limit of detection (LOD) for *P. clarkii* tissue in distilled water. Ten replicates per concentration. CCT is a conversion value of the threshold cycle of qPCR detection (CCT = 1000 / threshold cycle (cT)), where cT is the critical threshold number of qPCR reactions required for positive detection. A higher CCT indicates a higher eDNA concentration.

### Aquarium experiment

Each of the 21 aquarium samples was tested with three qPCR replicates, and 48 of the 63 reactions were positive. We successfully detected *P. clarkii* at densities from 100,000 crayfish / average paddy (9/9 replicates) down to just 1 crayfish / average paddy, but not perfectly (5/9 replicates, 55.6%, per aquarium: 2/3, 3/3, 0/3). We did not detect *P. clarkii* at the lowest tested density of 0.1 crayfish / average paddy (0/9 tests). The logarithm of CCT has a linear relationship with the logarithm of simulated crayfish density (linear regression, y = 1.40 + 0.02x, n = 63, R^2^ = 0.76, p < 0.001). We also carried out a mixed-effects model with village as a random factor, and the relationship remained highly significant at p < 0.001. There was an observable nonlinearity at the lowest densities (Fig 3).

**Fig 3.**
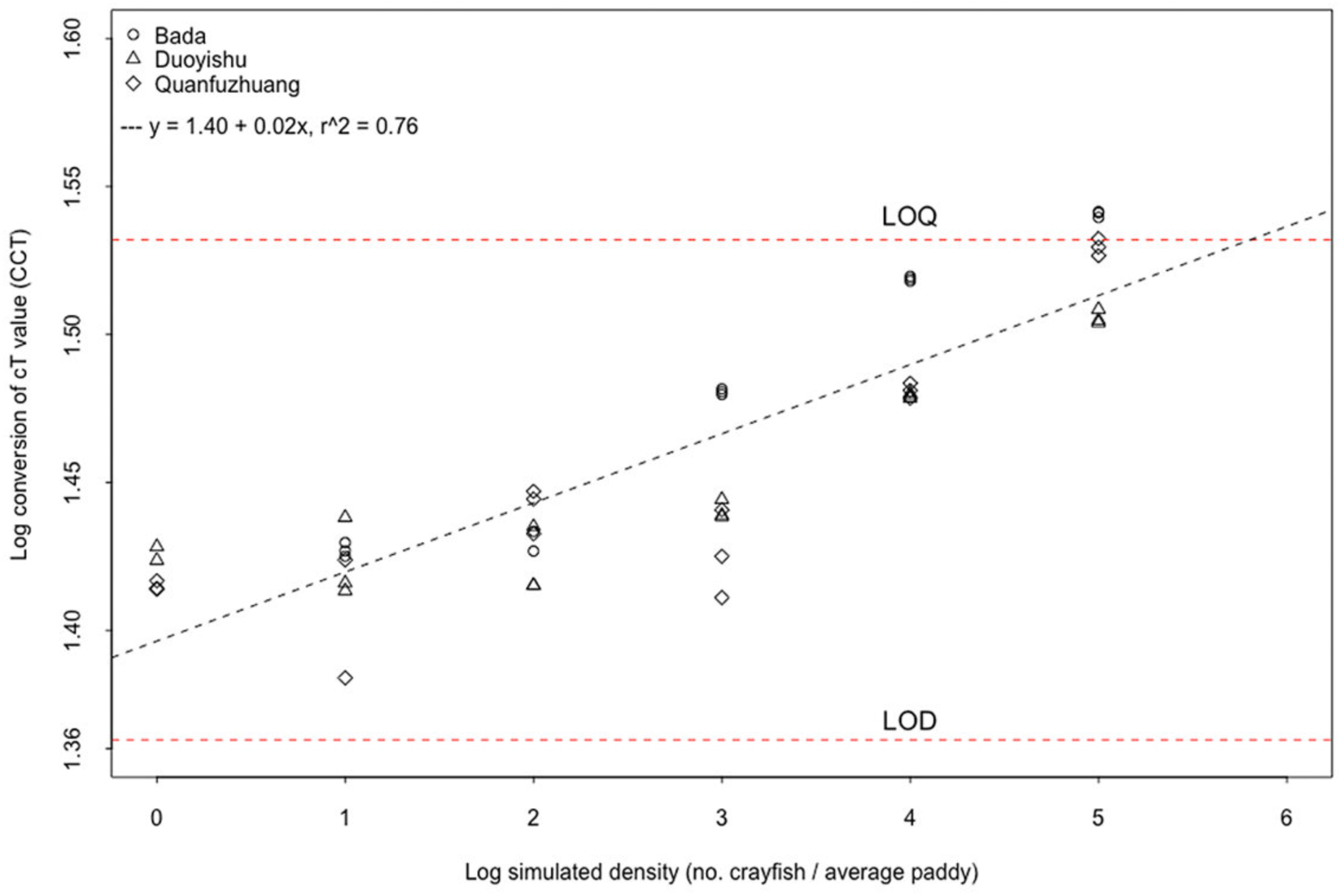
Environmental DNA detection and quantification in the aquarium experiment. The simulated density gradient of crayfish was achieved via a dilution series, where the average paddy has 10^5^ L of water. Logarithmic data has a better linear relation. Logarithm of CCT increases significantly with logarithm of crayfish density in all three sample sources. The estimated concentrations range lie between the Limit of Detection and the Limit of Quantification, as determined in Fig 2.

However, the range of CCT lay between the LOQ and LOD thresholds (Fig 2), so while we expect to be able to detect crayfish eDNA, we should expect that these CCT values are noisy estimates of eDNA concentrations. All DNA extraction blanks, and negative PCR controls were negative, indicating that contamination had not occurred.

### Field sampling and surveys

Of the 32 rice paddies that we sampled, farmers reported that 25 currently contained crayfish, with 18 “high” density and 7 “medium” density. Farmers did not report any “low” density paddies. eDNA successfully detected crayfish in all 25 crayfish-containing paddies, but bottle traps caught crayfish in only 17 of them (68% detection rate). Neither eDNA nor bottle traps detected crayfish in the 7 paddies where farmers and direct counting also did not report or detect crayfish. The qPCR inhibition test for these 7 crayfish-free negative samples indicated that inhibition is unlikely to be the reason that caused negative reactions. All DNA extraction blanks and PCR negative controls were negative.

Of the 25 paddies that had crayfish, after pesticide application, four were immediately invaded by large numbers of domestic ducks foraging for the exposed crayfish, which prevented us from carrying out direct counts. Nonetheless, we were able to census the remaining 21 crayfish-positive paddies and 7 crayfish-negative paddies for subsequent analyses. eDNA concentration predicts the direct count estimates more accurately (Fig 4A) than does the bottle-trap index (Fig 4B).

**Fig 4.**
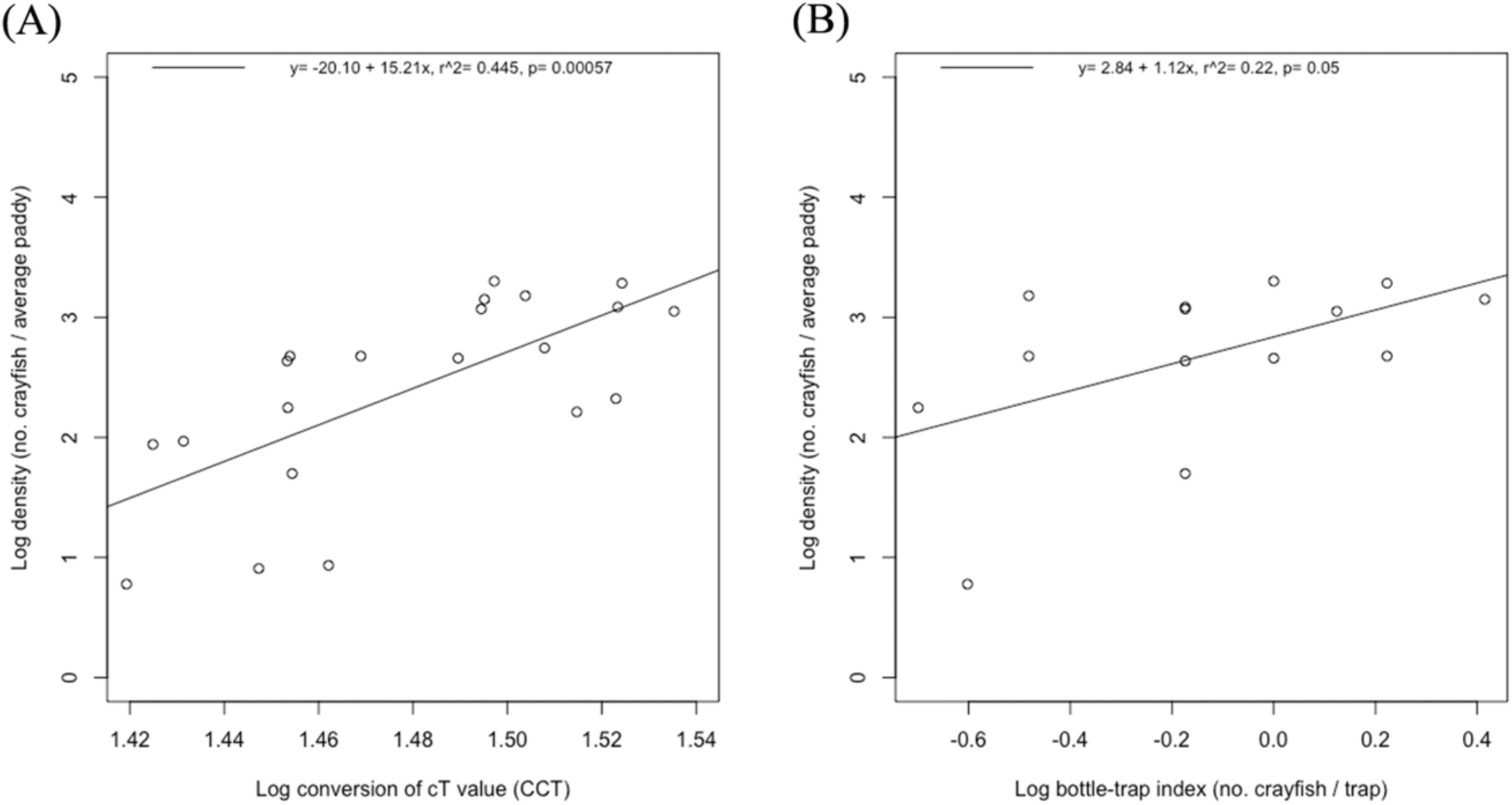
Comparison of eDNA and bottle-trapping with direct counts of crayfish. Direct counts are normalized to number of crayfish per average paddy (10^5^ L water). (A) the linear regression model for eDNA and direct counts of crayfish, and (B) for bottle-trapping and direct counts of crayfish.

We summarize our results in Figure 5, comparing farmer reports with estimated densities from eDNA, bottle-trap sampling, and direct counts, and we map the sampling locations (Fig 6) and show that the location with higher paddy densities (Qingkou) has no crayfish-free paddies, but the other two sites with lower densities have crayfish-free paddies.

**Fig 5.**
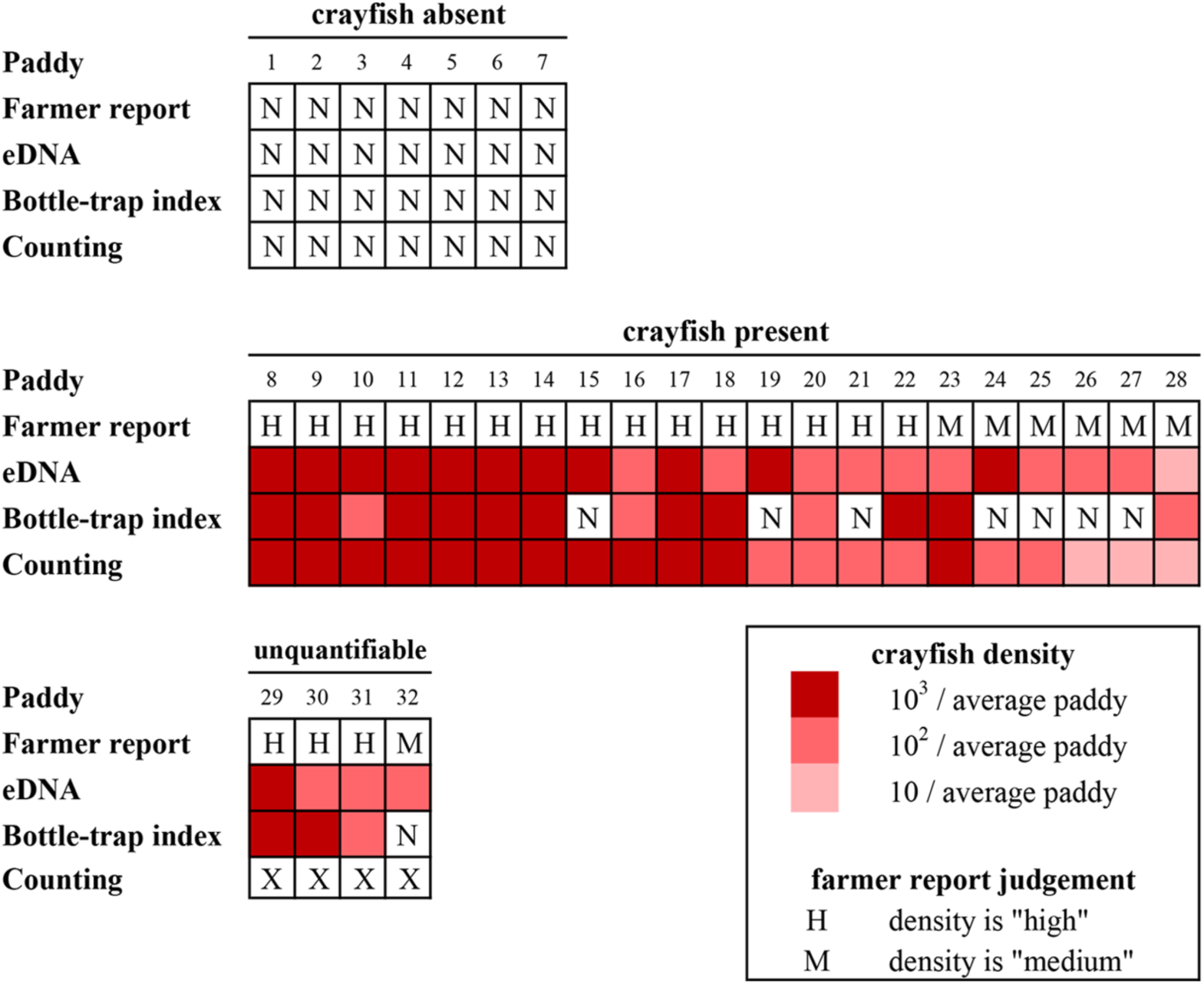
Comparison of farmer reports, eDNA, bottle-trap index, and direct counts. The 32 paddies are divided into seven that had no crayfish reported, 21 that had crayfish and could be checked with direct counting, and four that had crayfish but could not be checked with direct counting. (a) eDNA, bottle-traps, and direct counts all confirmed that the seven farmer-reported crayfish-free paddies were free of crayfish. (b) eDNA and direct counts confirmed that the 21 farmer-reported crayfish-containing paddies had crayfish, but bottle-traps failed to detect crayfish in 7 of those paddies, including one (paddy 15) that was deemed high density by farmer report, eDNA, and direct counts. (c) Finally, eDNA confirmed that all four uncounted farmer-reported, crayfish-containing paddies had crayfish, but bottle-traps failed to detect crayfish in one of them. The intensity of the red color indicates crayfish density estimates by direct counts, eDNA, and bottle trapping. The latter two were converted using the fitted models in Figure 4. N indicates no crayfish reported or detected. X indicates direct count not carried out. H and M indicate farmer reports of “high” and “medium” crayfish densities.

**Fig 6.**
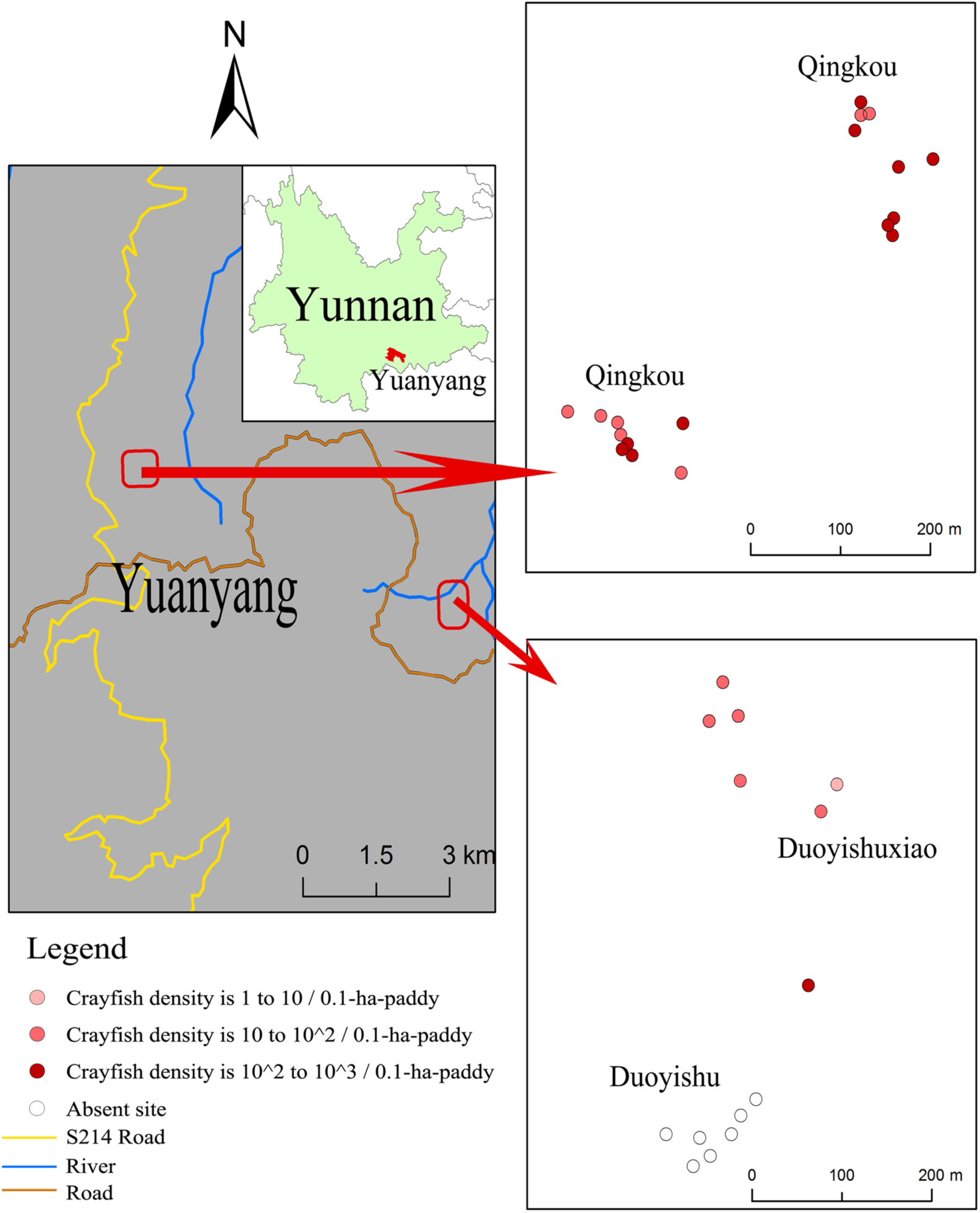
Map of sampling locations. Dots from dark red to light red are scaled for estimated crayfish density as in Figure 5

## Discussion

Our objective was to test whether eDNA could be used to detect invasive crayfish in the Honghe Hani rice terrace landscape, particularly at low densities. Reliable detection of absence could reduce use of pesticides and reliable detection of small populations could be used to detect invasion fronts or to adjudge successful eradication efforts.

We found that a single sampling event is enough to detect crayfish eDNA in all paddies where crayfish were directly observed (Fig 5), which is consistent with one published study [16] but different from another study where eDNA was only detected in 59% of sampling events [15]. In contrast, bottle-traps were prone to false negatives, detecting crayfish in only 68% of crayfish-containing paddies (Fig 5). Traps therefore will require multiple site visits to reach the same efficiency as eDNA in this system, and they capture non-target organisms, such as tadpoles, fish fry, and aquatic invertebrates. Furthermore, eDNA does not appear to have resulted in any false positives (Fig 5), as determined by farmer reports and direct counts. At the quantitative level, by taking advantage of the fact that farmers in Yuanyang annually kill crayfish with pesticide before planting rice seedlings, we found that eDNA could estimate crayfish densities per paddy more accurately than could trap sampling (Fig 4), which suggests that it is not necessary to rely solely on un-normalized qualitative reports by farmers.

However, the aquarium results caution that at the lowest simulated density of 1 crayfish per average paddy, it is nonetheless possible to achieve a false-negative result (one tank returned 0/3 crayfish amplifications). The aquarium experiments also demonstrate that the eDNA estimates cannot be taken as very accurate, as realistic eDNA concentrations are below the Limit of Quantification (Fig. 3), and this caveat is underlined by the less than perfect correlation between the quantity estimates from eDNA and direct counts (Fig 4A).

Also, we caution that our aquaria contained only water from the paddies, while real paddies also have a layer of watery mud below the liquid water layer. Given that the crayfish burrow into this mud and into the paddy walls, it is possible that some, perhaps even a large amount, of crayfish eDNA is *directly* injected into the mud and thus never available for water sampling [31]. We could not detect crayfish at a simulated density of 0.1 crayfish per average paddy, which implies that if a large proportion of crayfish eDNA does directly go into the mud, very small populations are likely not to be detected.

Taken together, this possibility of false negatives due to eDNA loss to mud and less-than-perfect qPCR detection at the lowest densities, counterbalanced by the observed high rate of true-positive detections and low rate of false-positive detections in our water samples (Fig 5) supports the use of occupancy modeling to model the distribution of crayfish, as has been suggested for eDNA detection of chytrid fungus from water samples [32], thereby identifying portions of the landscape where crayfish are still absent and where surveillance efforts would be most efficiently targeted. An example of this is Duoyishu village, where all but one of the surveyed paddies were free of crayfish (Fig 6, hollow dots), but the paddies to the north represent an invasion front. Potentially, digital droplet PCR, which is more sensitive to eDNA [33], in part because it is less affected by PCR inhibition, might be a more effective method for detecting very low concentrations of crayfish eDNA.

In this study, we did not attempt to correct for environmental variation (e.g. pH, temperature, solar radiation), but crayfish may vary considerably in their release rate of DNA [34-37]. Food availability directly governs excretion rates, and juvenile crayfish molt frequently, but adults do not, and such exuviae are a known source of eDNA [38]. It is possible that eDNA detection will be more or less likely at certain times of the year, and it will be necessary to test for this in the future. The advantage of the Honghe Hani system is that it is possible to validate eDNA results against direct counting, which is rarely possible to do with water-borne eDNA.

In conclusion, eDNA sampling is clearly more efficient than bottle-trap-sampling in this system, given that the primer/probe set is already developed and that it is acceptable to deem crayfish present with less than perfect eDNA detection (given the alternatives now of farmer self-reporting and repeated bottle-trapping) [39]. It is now feasible to trial eDNA sampling to detect new invasion fronts, to assess the need for insecticide applications in low-crayfish zones, and to inform any eventual eradication campaign.

## Acknowledgments

We thank the South China Barcode Center for logistical support; Yunnan Normal University for field assistance; and Yuanyang Agricultural Administrative Department for assistance with interviews.

## Supporting Information

**Zip S1** The R running script and the Excel file used for all data analyses

